# ALK1-BMPRII agonism by clustering bispecific antibodies treats hereditary hemorrhagic telangiectasia

**DOI:** 10.1101/2025.08.13.670104

**Authors:** Sima Qutaina, Haitian Zhao, Zhiming Wang, Andy Sullivan, Melissa Geddie, Raminderjeet Kaur, Hong Tian, Huan Yue, Xin Wang, Siyang Guo, Margherita Bruni, Erica Christen, Fabien Campagne, Helen M. Arthur, Patrick André, Philippe Marambaud

**Affiliations:** Litwin-Zucker Alzheimer Research Center and Institute of Molecular Medicine, The Feinstein Institutes for Medical Research, Northwell Health, Manhasset, NY, USA; Diagonal Therapeutics, Watertown, MA, USA; Northwell, New Hyde Park, NY and Otolaryngology and Facial Plastics, Great Neck, NY, USA; Campagne Machine Intelligence Consulting LLC, New York, NY, USA; Biosciences Institute, Newcastle University, Newcastle, UK; Donald and Barbara Zucker School of Medicine at Hofstra/Northwell, Hempstead, NY, USA

## Abstract

Hereditary hemorrhagic telangiectasia (HHT) is characterized by arteriovenous malformations (AVMs) and severe bleeding caused by loss-of-function mutations in the ALK1 receptor pathway. We developed clustering agonist bispecific antibodies (BsAbs) targeting ALK1 and its activating partner, the Ser/Thr receptor kinase BMPRII. These BsAbs induced ALK1-BMPRII proximity association, stimulated the downstream Smad1/5/8 signaling cascade, and treated HHT pathologies in various mouse models. BsAb treatment reduced AVM burden by up to 95% in HHT mice, preventing anemia, cardiomegaly, and premature death. The BsAbs also enhanced Smad1/5/8 signaling in endothelial cells derived from HHT patients with ALK1 mutations and prevented retinal AVMs in a newly developed knock-in mouse carrying an HHT-causing ALK1 mutation. These findings establish ALK1-BMPRII agonism as a promising therapeutic strategy for HHT.

## Introduction

Members of the TGF-β superfamily of ligands and receptors function as biological sentinels, maintaining vascular quiescence throughout the entire vascular tree (*1*). Heterozygous germline loss-of-function (LoF) mutations in the endothelial ENG-ALK1-Smad4 axis—a TGF-β signaling pathway—result in the vascular dysplasia termed hereditary hemorrhagic telangiectasia (HHT) (*2*). Most patients carry mutations in either *ENG* [encoding endoglin (ENG)] or *ACVRL1* [encoding activin receptor-like kinase 1 (ALK1)], which result in HHT type 1 (HHT1) or type 2 (HHT2), respectively. Additionally, rare mutations in *SMAD4* [encoding small mothers against decapentaplegic homolog 4 (Smad4)] cause juvenile polyposis-HHT combined syndrome (JP-HHT) (*3*).

HHT affects approximately one in 5,000 individuals and is characterized by the focal development of arteriovenous malformations (AVMs) in various organs and tissues. AVMs result from abnormal enlargements of blood capillaries, leading to fragile, hemorrhage-prone, high-flow shunts between arteries and veins (*4*). Depending on which vascular beds are affected, HHT can manifest as epistaxis or hemorrhage in the gastrointestinal tract, liver, lungs, or brain, causing severe anemia in about 60% of the patient population. HHT can also raise the risk of stroke, organ failure, and cardiac complications (*5, 6*).

ENG functions as a blood flow-dependent auxiliary receptor, presenting BMP9 and BMP10 ligands to signaling receptor complexes composed of the BMP type I receptor ALK1 and a BMP type II receptor, such as BMPRII (*7*–*9*). This interaction enables the constitutively active Ser/Thr receptor kinase BMPRII to activate ALK1 through physical proximity and phosphorylation of ALK1’s GS domain (*10, 11*). ALK1 subsequently induces the phosphorylation of signal transducers Smad1, 5, and 8, leading to the recruitment of Smad4 into Smad1/5/8-Smad4 transcriptional nuclear complexes that regulate specific gene expression programs involved in vascular quiescence (*12*–*14*). Despite recent progress in understanding the biological processes that drive receptor activation and signal transduction, there is limited evidence to suggest the feasibility of correcting the root cause of the disease (*15, 16*), namely, the defective receptor signaling machinery in HHT1 and HHT2.

In this study, we introduce ALK1-BMPRII clustering bispecific antibodies (BsAbs) and demonstrate their ability to activate ALK1 signaling and inhibit AVM pathologies in BMP9/10- and ENG-deficient mice, as well as in heterozygous knock-in (KI) mice carrying a LoF HHT2 mutation. ALK1-BMPRII BsAbs also showed ALK1-activating properties in blood-outgrowth endothelial cells (BOECs) from HHT2 patients. This study demonstrates that targeted ALK1-BMPRII agonism by BsAbs can effectively enhance ALK1 signaling to block HHT pathogenesis.

## Results

### ALK1-BMPRII BsAbs promote ALK1 and BMPRII association and activate ALK1 signaling

Using high-throughput screening of repertoires and proprietary computational analyses, we identified BsAbs with high specificity for the extracellular domains of ALK1 and BMPRII. Antibody linker optimization, which involved adjusting the linker’s length and flexibility to promote the proper orientation and function of the BsAbs, was used to select candidate antibodies. We selected the dual variable domain immunoglobulin (DVD-Ig) BsAb DIAG100 due to its high *in vitro* affinity and dual engagement with both human and mouse recombinant ALK1 and BMPRII (Figs. 1A, 1B). A cell-based protein interaction assay (DiscoverX PathHunter) showed that DIAG100 has potent ALK1-BMPRII clustering capabilities (Fig. 1C). Western blot (WB) analyses were then conducted on protein extracts of mouse endothelial Mile Sven 1 (MS1) cells incubated in serum-depleted medium and treated for 24h with increasing doses of DIAG100. An elevation of phospho-Smad1/5/8 (p-Smad1/5/8) and its transcriptional target, ID3, was detected at doses of 300 ng/mL DIAG100 and above, indicating that DIAG100, like BMP9, can activate ALK1 signaling under these culture conditions (Fig. 1D). 300 ng/mL DIAG100 also significantly increased p-Smad1/5/8 and ID3 levels in MS1 cells incubated in growth factor-containing medium (Figs. 1E, 1F), a more physiologically relevant condition and an important prerequisite for the *in vivo* evaluation of the BsAbs. In addition, DIAG100 demonstrated sustained activation of p-Smad1/5/8 and ID3 for at least 72h in MS1 cells incubated in growth factor-containing medium (Figs. 1G, 1H).

**Figure 1:**
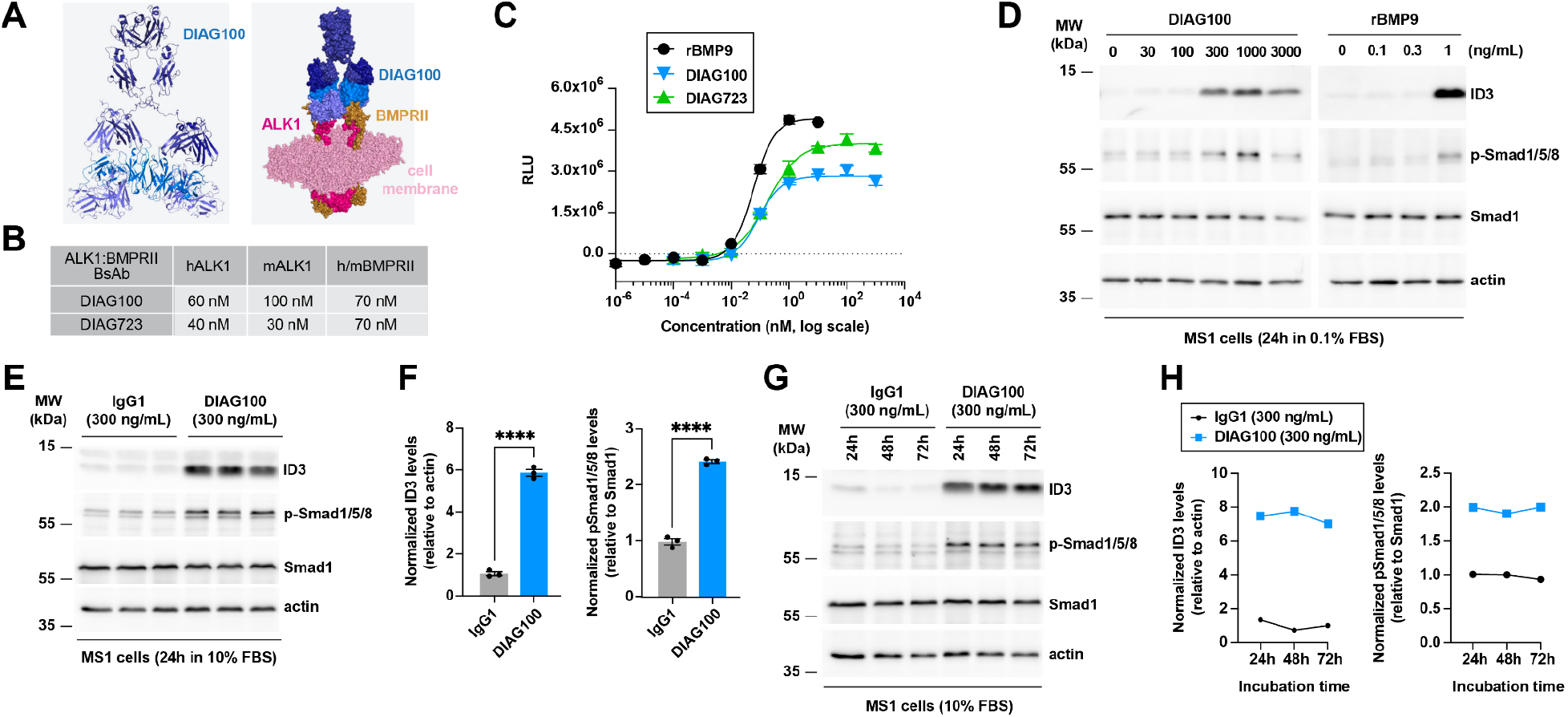
ALK1-BMPRII BsAbs promote ALK1 and BMPRII association and activate ALK1 signaling. (**A**) DIAG100 structure and computational modeling of its interaction with ALK1 and BMPRII at the cell membrane. (**B**) DIAG100 and DIAG723 affinities for human and mouse recombinant ALK1 and BMPRII. (**C**) DiscoverX assay measuring ALK1 and BMPRII proximity association by DIAG100, DIAG723, and recombinant BMP9 (rBMP9). RLU, relative light units. (**D**) WB analyses of the indicated proteins from MS1 cells treated with different concentrations of DIAG100 or rBMP9 in serum-depleted medium (0.1% FBS) for 24h. (**E**) WB analyses of the indicated proteins from MS1 cells treated with 300 ng/mL DIAG100 or IgG1 control in complete medium (10% FBS) for 24h. (**F**) Quantification of the levels of ID3 and p-Smad1/5/8 from the WB analysis in (E) (n = 3). Data are presented as mean ± s.e.m., unpaired t-test. ****p < 0.0001. (**G**) WB analyses of the indicated proteins from MS1 cells treated with 300 ng/mL DIAG100 or IgG1 control in complete medium (10% FBS) for the indicated times. (**H**) Quantification of the levels of ID3 and p-Smad1/5/8 from the WB analysis in (G).

### DIAG100 binds to the retinal endothelium and promotes ALK1-BMPRII clustering in vivo

DIAG100 was labeled with Alexa Fluor 647 (AF647) to track its localization in the retinal vasculature after injection into the bloodstream. DIAG100-AF647—which retained its clustering capabilities and Smad1/5/8 signaling activation properties (Fig. S1)—or IgG1-AF647 (used as a control) was injected intracardially into anesthetized postnatal day 7 (P7) mice. Five minutes post-injection, the blood was flushed with PBS to remove unbound labeled antibodies from circulation. Immunofluorescence (IF) analyses showed that DIAG100-AF647 strongly decorated the retinal vasculature by binding to the endothelium of arteries, capillaries, and veins, while IgG1-AF647 did not (Figs. 2A, 2B). Additionally, DIAG100-AF647 strongly bound to the endothelium of developing vessels at the front of the veins (Fig. 2A), an area of the retinal vasculature that is already lumenized and contains blood (*17*). DIAG100-AF647 also effectively bound to the endothelium of the AVMs in the retina of mouse neonates treated with anti-BMP9 and anti-BMP10 blocking antibodies, hereafter referred to as the BMP9/10 immunoblocked (BMP9/10ib) mice (Fig. 2C), a model of HHT that has been extensively characterized in our lab (*15, 18*–*20*). A proximity ligation assay (PLA) assessing ALK1 and BMPRII interaction revealed a robust and significant elevation of PLA signals on arterial retinal ECs following DIAG100 intraperitoneal (i.p.) injection in mice (Fig. 2D, 2E). Thus, the BsAb administered either intracardially or intraperitoneally enters the bloodstream, binds to the endothelium, and promotes endothelial ALK1-BMPRII clustering *in vivo*.

**Figure 2:**
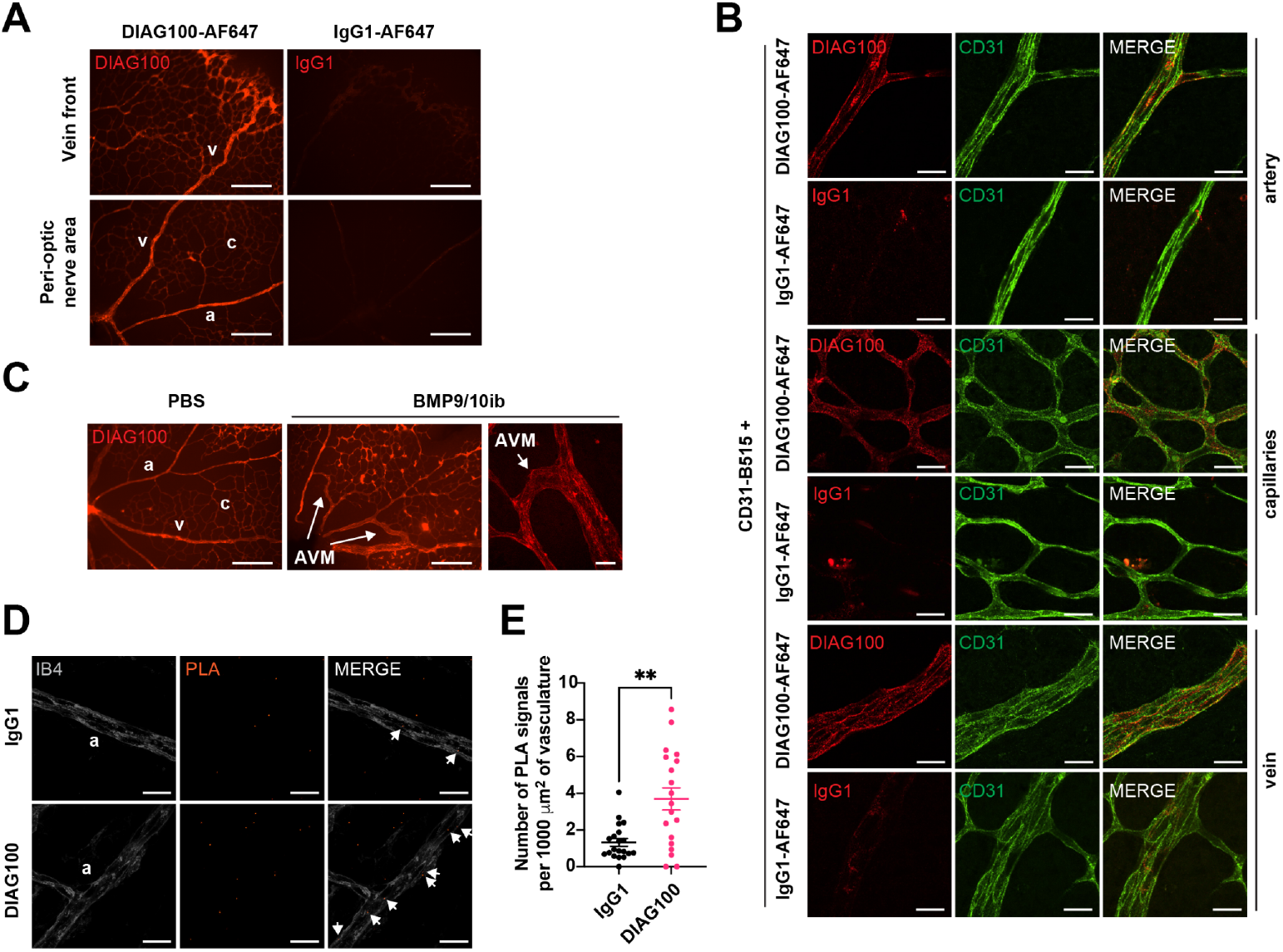
DIAG100 binds to the retinal endothelium and promotes ALK1-BMPRII clustering *in vivo*. (**A**) Representative images of retinas from P7 C57BL/6 mice stained with intracardially injected DIAG100-AF647 or IgG1-AF647 control (3 mg/kg). Areas at the front of a vein (upper panels) and around the optic nerve (lower panels) are shown. Scale bars, 200 µm. (**B**) Representative images of retinas from P7 C57BL/6 mice stained with intracardially injected CD31-B515, and DIAG100-AF647 or IgG1-AF647 control (1 mg/kg), showing arteries, capillaries, and veins. Scale bars, 20 µm. (**C**) Representative images of retinas from P7 PBS control and BMP9/10ib mice stained with intracardially injected DIAG100-AF647 (3 mg/kg). Scale bars, 200 µm (left two panels), 50 µm (right panel). (**D**) Representative images of retinas from P7 C57BL/6 mice injected i.p. on P6 with DIAG100 or IgG1 control (3 mg/kg) stained with isolectin B4 (IB4, white) and analyzed with ALK1-BMPRII PLA. Arrows denote PLA signals in ECs (IB4^+^ cells). Scale bars, 20 µm. (**E**) Quantification of the PLA signals visualized as in (D). Data are presented as mean ± s.e.m., unpaired t-test with Welch’s correction. **p < 0.01. a, artery; c, capillaries; v, vein.

### DIAG100 treats AVMs in BMP9/10ib and EngiECKO mice

Our observations that DIAG100: (i) bound to ALK1 and BMPRII, (ii) promoted ALK1-BMPRII association *in vitro* and *in vivo*, (iii) bound *in vivo* to the endothelium, and (iv) activated ALK1 signaling in ECs led us to evaluate its anti-AVM properties in HHT models. Following disease induction by i.p. injection of BMP9 and BMP10 blocking antibodies on P3 and P4 [BMP9/10ib model (*20*), Fig. 3A], retinas developed prominent AVMs around the optic nerve (Fig. 3B, white arrows). A dose-response showed that treatment with 1 mg/kg DIAG100 on P3+P4 (i.p.) led to a nearly complete prevention of retinal AVM development (95% reduction on average, p < 0.0001, ordinary one-way ANOVA, Dunnett’s multiple comparisons test; Figs. 3B, 3C). DIAG100 did not cause any apparent toxicity in the pups, and body weight remained unaffected at the end of the experiment (Fig. 3C). We then asked whether DIAG100 could ameliorate preexisting AVMs using a ‘regression protocol’ that involves treating BMP9/10ib mice starting on P6 [Fig. 3D and Refs. (*19, 20*)], a time when AVMs are already formed (Fig. 3E, left images). Treatment with 1 mg/kg DIAG100 on P6+P7 (Fig. 3D) resulted in nearly complete regression by P8 of AVMs that developed on P6 (Figs. 3E, 3F). Thus, DIAG100 can both prevent and reverse AVMs in the retina of the neonatal BMP9/10ib model.

**Figure 3:**
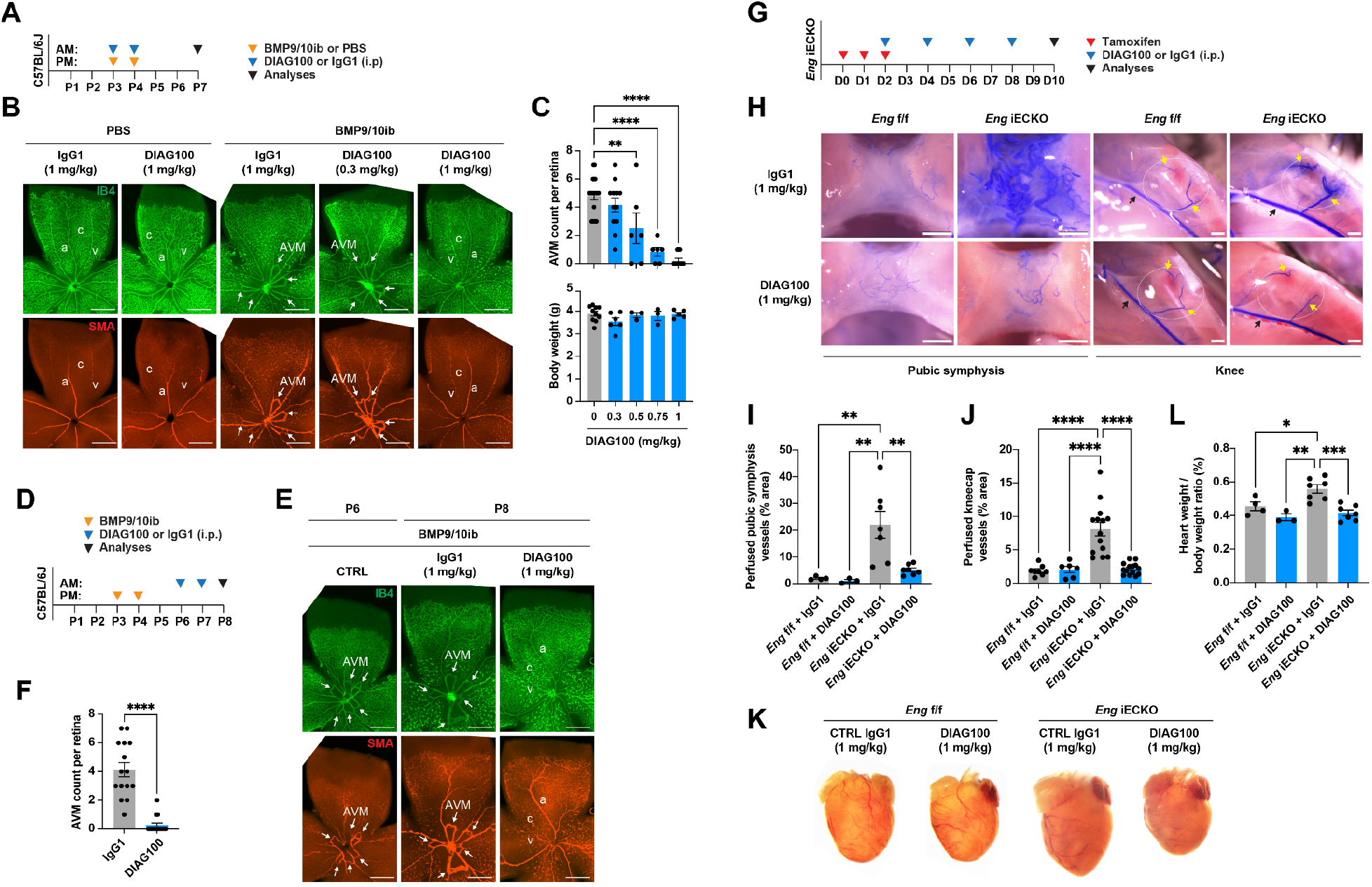
DIAG100 treats AVMs in BMP9/10ib and *Eng*^iECKO^ mice. (**A**) Schematic representation of the injection protocol used in (B) and (C). AM, morning; PM, afternoon. (**B**) Representative staining with isolectin B4 (IB4, green) and of α-smooth muscle actin (SMA, red) in whole-mount retinas from PBS control and BMP9/10ib mice treated (i.p.) with the indicated doses of DIAG100 or IgG1 control. (**C**) The upper bar graph shows the number of retinal AVMs and the lower bar graph shows the body weight of BMP9/10ib mice treated as in (B) with the indicated doses of DIAG100. Data are presented as mean ± s.e.m., one-way ANOVA with Tukey’s multiple comparisons test. (**D**) Schematic representation of the injection protocol used in (E) and (F). (**E**) Representative staining with IB4 (green) and of SMA (red) in whole-mount retinas on P6 (left panels) and P8 (middle and right panels) from BMP9/10ib mice, untreated (control, CTRL) or treated i.p. with IgG1 or DIAG100 (1 mg/kg). (**F**) The bar graph shows the number of retinal AVMs from BMP9/10ib mice treated as in (E). Data are presented as mean ± s.e.m., unpaired t-test. Both eyes of each mouse were analyzed in (C) and (F). a, artery; c, capillaries; v, vein. Arrows denote AVMs. Scale bars, 500 µm (B, E). (**G**) Schematic representation of the experimental layout used in (H-L). (**H**) Representative bright-field images of pubic symphyses and knees from *Eng*^iECKO^ mice and *Eng*^f/f^ control littermates perfused with blue latex beads and treated i.p. with IgG1 or DIAG100 (1 mg/kg). The black arrows denote the descending genicular artery, while the yellow arrows denote the superior and inferior medial genicular arteries. Scale bars, 1 mm. (**I, J**) The bar graphs show the area occupied by latex-perfused vessels as a % of the total pubic symphysis area (I) and total kneecap area (J, marked by the circles) from *Eng*^f/f^ and *Eng*^iECKO^ mice treated as in (G). Data are presented as mean ± s.e.m., one-way ANOVA with Tukey’s multiple comparisons test. (**K, L**) Representative mouse hearts (K) and heart weight/body weight ratio measurements (L) from *Eng*^f/f^ and *Eng*^iECKO^ mice treated with IgG1 or DIAG100 as in (G). Data are presented as mean ± s.e.m., one-way ANOVA with Tukey’s multiple comparisons test. *p< 0.05, **p < 0.01, ***p < 0.001, ****p < 0.0001 (C, F, I, J, L).

To determine the effect of ALK1-BMPRII activation in a different HHT model and beyond the neonatal stage, we used *Eng*iECKO mice (*21, 22*). Evidence shows that ten days after gene deletion induction, *Eng*iECKO mice develop prominent AVMs in the pubic symphysis, which can be visualized using blue latex dye injections (*23*). In 8-week-old homozygous *Eng*iECKO mice, we confirmed significant development of pubic symphysis AVMs compared to *Eng*f/f control littermates (Figs. 3G, 3H). Treatment with 1 mg/kg DIAG100 on Days 2, 4, 6, and 8 after gene deletion nearly completely prevented AVM development by Day 10, while treatment with 1 mg/kg IgG1 control had no effect (Figs. 3G-3I). Further investigation revealed that *Eng*iECKO mice also develop surface vascular defects around the knees (Fig. 3H). The main arteries projecting above and below the kneecap are the superior and inferior medial genicular arteries. Blue latex dye perfusion enabled visualization of the descending genicular artery (Fig. 3H, black arrows) and the medial genicular arteries (Fig. 3H, yellow arrows). In *Eng*iECKO mice, we observed dilation of the medial genicular arteries and the formation of vessel bundles at their extremities above the kneecap. Treatment with DIAG100 prevented the increase in vascularization of the kneecap surface (Figs. 3H, 3J). *Eng*iECKO mice also develop cardiomegaly, likely due to hemodynamic changes caused by peripheral AVMs (*23*). Treatment with 1 mg/kg DIAG100 prevented heart enlargement in *Eng*iECKO mice, whereas 1 mg/kg IgG1 control did not (Figs. 3K, 3L). Overall, these results demonstrate that DIAG100 both prevents and reverses AVMs under BMP9/10 and ENG LoF conditions, bypassing the regulation by both ligands and the co-receptor.

### DIAG100 activates ALK1 signaling in HHT2 patient BOECs and improves AVM pathology in HHT2 KI mice

An important question is whether the ALK1-BMPRII BsAb can activate ALK1 signaling in the presence of HHT mutations in ALK1. We previously reported the generation of BOECs isolated from HHT patients and healthy controls (*15, 19*). BOECs from one healthy donor and two HHT2 patients with mutations in the ALK1 extracellular domain [ALK1-G48E (*24*)] and kinase domain [ALK1-T372fs* (*25*)] were tested. 1 μg/mL DIAG100 significantly increased levels of p-Smad1/5/8, ID3, and ID1 (another Smad4 transcriptional target) in both healthy donor and HHT2 BOECs (Figs. 4A, 4B). To better mimic human HHT genetics, we generated a KI mouse carrying the well-characterized HHT2-causing and kinase-dead ALK1-R479* mutant [*Acvrl1*R479* (*26, 27*)]. Heterozygous *Acvrl1*R479*/+ mice were viable and fertile, and examination of their retinas at the neonatal stage revealed no obvious vascular defects (Fig. S2). As expected, treating *Acvrl1*R479*/+ pups and their *Acvrl1*+/+ littermates with BMP9/10 blocking antibodies induced retinal AVMs (Figs. 4C-4E). 1 mg/kg DIAG100 significantly prevented AVMs in both *Acvrl1*R479*/+ and *Acvrl1*+/+ pups (Figs. 4C-4E). Overall, these findings demonstrate that DIAG100 can activate ALK1 signaling and improve AVM pathology in the presence of germline ALK1 mutations associated with HHT.

**Figure 4:**
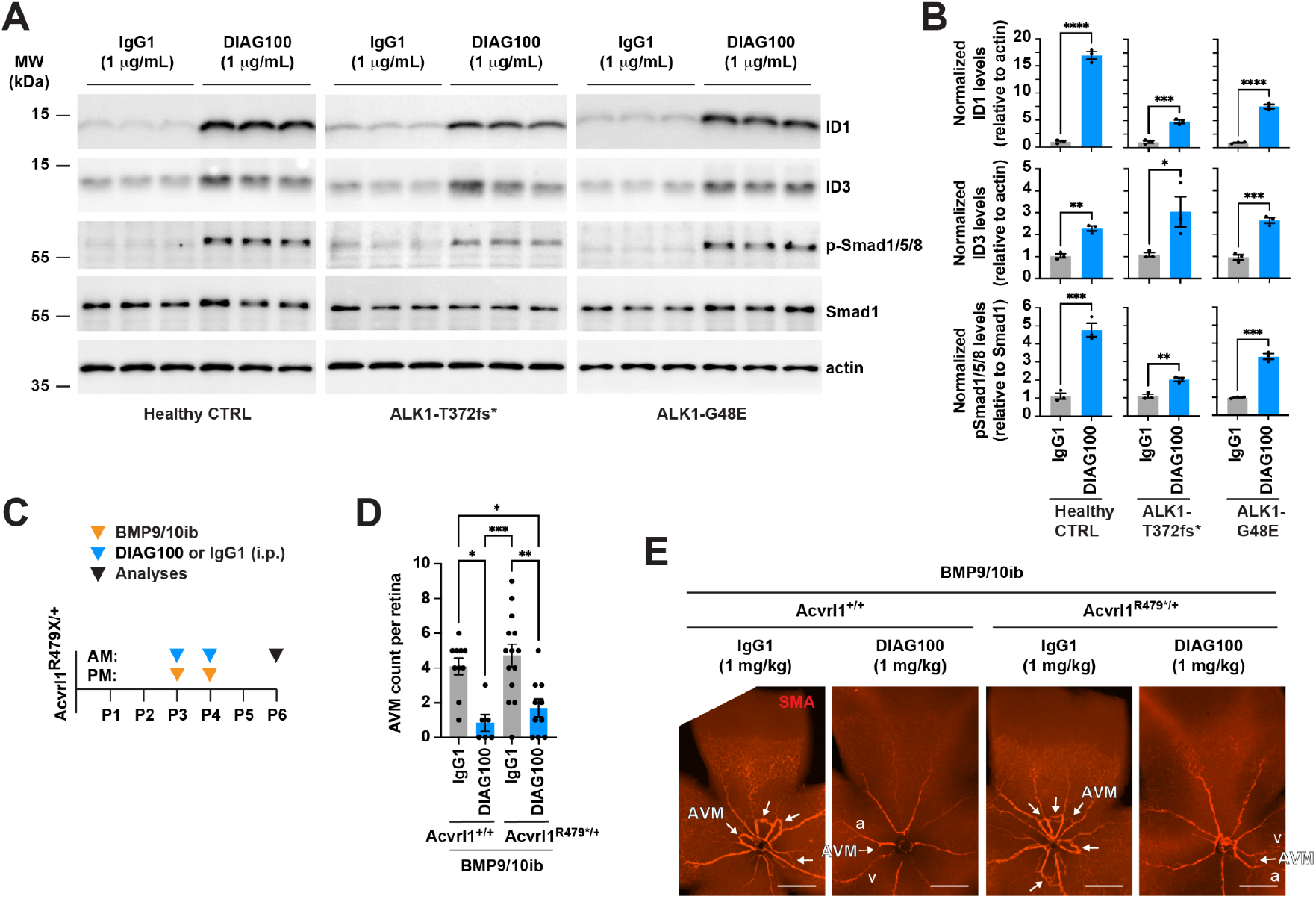
DIAG100 activates ALK1 signaling in HHT2 patient BOECs and improves AVM pathology in HHT2 KI mice. (**A**) WB analyses of the indicated proteins from healthy donor and HHT2 patient BOECs serum-depleted for 3h (0.1% FBS) and then treated with 1 μg/mL DIAG100 or IgG1 control for 2h. (**B**) Quantification of the levels of ID1, ID3, and p-Smad1/5/8 from the WB analysis in (A) (n = 3). Data are presented as mean ± s.e.m., unpaired t-test. (**C**) Schematic representation of the injection protocol used in (D, E). (**D, E**) The bar graph shows the number of retinal AVMs (D) and representative staining of SMA in whole-mount retinas from BMP9/10ib;*Acvrl1*^R479*/+^ mice and BMP9/10ib;*Acvrl1*^+/+^ controls injected (i.p.) with DIAG100 or IgG1 control (1 mg/kg). a, artery; v, vein. Arrows denote AVMs. Scale bars, 500 µm. Data in (D) are presented as mean ± s.e.m., one-way ANOVA with Tukey’s multiple comparisons test. *p< 0.05, **p < 0.01, ***p < 0.001, ****p < 0.0001 (B, D).

### ALK1-BMPRII receptor controls a subset of BMP9-regulated genes

ALK1-BMPRII agonist BsAbs can serve as a unique tool for studying the specific signaling response of the ALK1-BMPRII receptor complex. We interrogated and compared the transcriptomes of human umbilical vein endothelial cells (HUVECs) treated with DIAG100 or BMP9 to determine the DIAG100 gene expression signature and how it compares to the BMP9 gene expression signature. Gene expression deregulation was observed in both treatment groups (Figs. 5A, 5B), and a linear correlation was found when comparing their respective log2 fold changes (Fig. 5C). We identified 69 genes significantly deregulated by DIAG100 treatment [false discovery rate (FDR) adjusted p-value ≤ 5%, log2 fold change (FC) cutoff = 0.85]. All 69 of these genes belonged to the group of BMP9-regulated genes (Fig. 5D), showing that DIAG100 did not produce off-target transcriptional responses, at least *in vitro*, as no genes outside BMP9 control were found to be deregulated by the BsAb. Within these 69 genes, two groups were identified (Fig. 5E): a group of 35 genes (group #1) that were deregulated (either up or down) with an amplitude that was not significantly different between DIAG100 and BMP9 treatments (Fig. 5F), and a group of 34 genes (group #2) that were deregulated by DIAG100 but with a significantly weaker amplitude compared to BMP9 (Fig. 5G). ID1 and ID3 were part of group #1 (Fig. 5F), and WB analysis confirmed that DIAG100 and BMP9 increased their expression with a similar amplitude (Fig. 5H). SGK1 was in group #2, and WB analysis confirmed that BMP9 was more potent than DIAG100 in increasing its expression (Fig. 5H). Thus, DIAG100-driven ALK1-BMPRII activation is associated with the regulation of a specific subset of BMP9-regulated genes.

**Figure 5:**
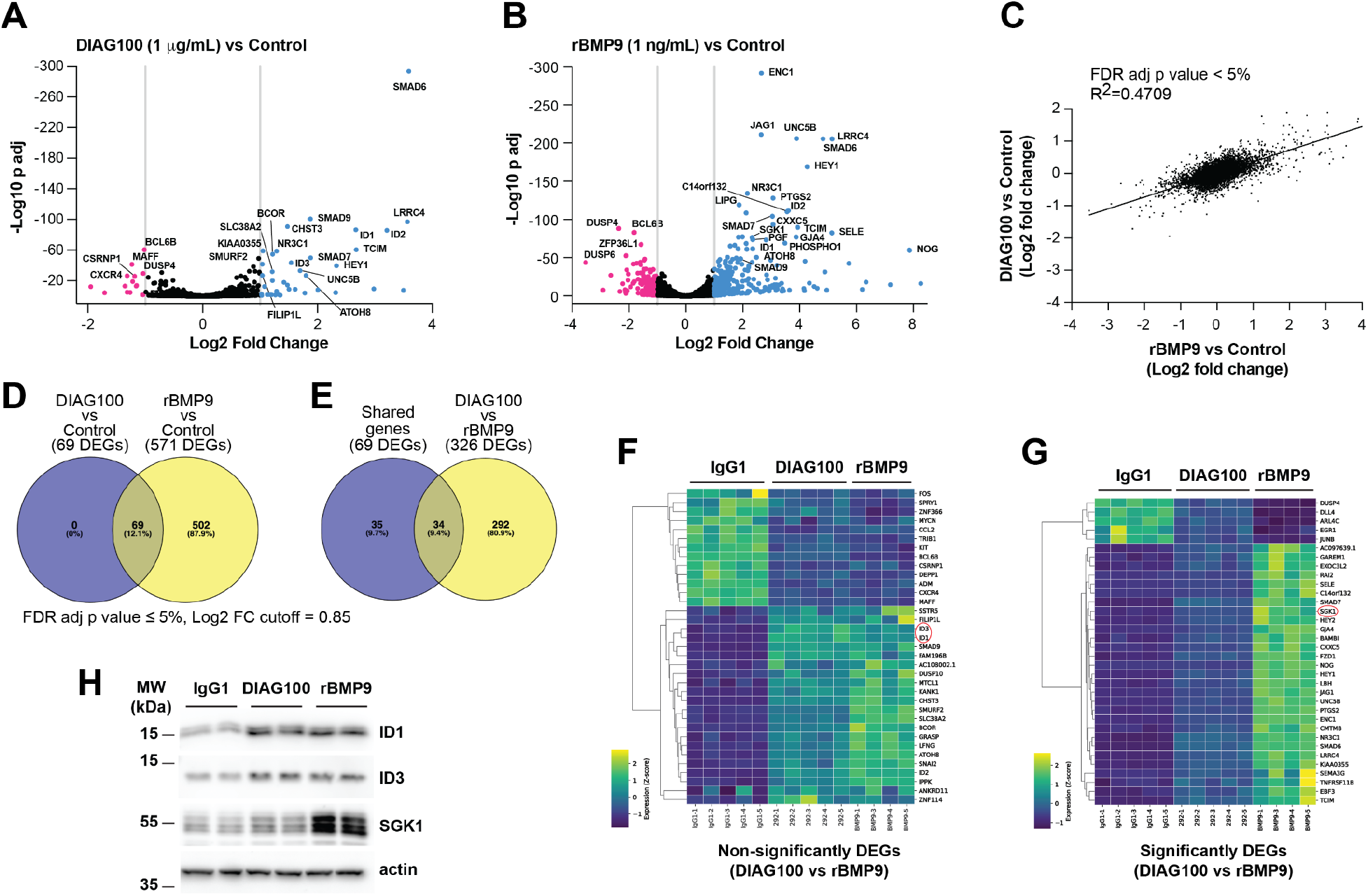
ALK1-BMPRII receptor controls a subset of BMP9-regulated genes. (**A, B**) Volcano plots showing differentially expressed genes (DEGs) in HUVECs serum-depleted for 2h (0.5% FBS) and then treated for 4h with 1 μg/mL DIAG100 [n=5 biological replicates, (A)] or 1 ng/mL rBMP9 [n=4, (B)], relative to 1 μg/mL IgG1 (Control, n=5). Representative significantly DEGs are identified. At the concentrations used, DIAG100 is applied at ∼60-fold molar excess compared to BMP9. (**C**) Log2 fold change slope comparing the DEGs in the DIAG100 vs control comparison to the DEGs in the rBMP9 vs control comparison. (**D**) Venn diagram depicting the relationship between the DEGs of the DIAG100 vs control comparison and the DEGs of the rBMP9 vs control comparison. (**E**) Venn diagram depicting the relationship between the shared DEGs of the comparison in (D) and the DEGs of the DIAG100 vs rBMP9 comparison. (**F, G**) Heatmaps showing the shared DEGs of the comparison in (D) that are either not significantly different (F) or significantly different (G) in the DIAG100 vs rBMP9 comparison. (**H**) WB analyses of the indicated proteins from HUVECs treated as in (A, B).

### DIAG723 blocks AVM development, anemia, and lethality in BMP9/10ib mice

Further optimization of DIAG100 was conducted to enhance its target engagement and clustering potency. DIAG723 (Fig. 1B) was identified as having improved ALK1-BMPRII clustering capabilities in the DiscoverX cell-based protein interaction assay compared to DIAG100 (Fig. 1C). Smad1 signaling activition by DIAG723 was demonstrated across several human endothelial cell lines [human pulmonary artery endothelial cells (HPAECs), telomerase immortalized microvascular endothelial cells (TIME), and human microvascular endothelial cells-1 (HMEC-1); Fig. S3] and proved more potent than DIAG100 at increasing p-Smad1/5/8 and ID3 levels in MS1 (Fig. 6A). When injected i.p. up to 1.5 mg/kg, DIAG723 significantly and dose-dependently prevented retinal AVMs in BMP9/10ib pups (Fig. S4). Pharmacokinetic studies in C57BL/6 mice showed that after a 1 mg/kg i.p. injection, DIAG723 reached a peak blood concentration of about 40 μg/mL at 8h post-injection, then declined rapidly. In contrast, when DIAG723 was administered subcutaneously (s.c., behind the neck) at 1 mg/kg, it also peaked in the blood at around 40 μg/mL at 8h but maintained this concentration for up to two days (Fig. 6B). We confirmed that 1 mg/kg DIAG723 administered s.c. effectively blocked retinal AVMs (88% reduction on average, p < 0.0001, unpaired t-test; Figs. 6C-6E). Exploiting the sustained delivery from s.c.injection, we performed longer-term treatments with DIAG723 and evaluated its effects on anemia and survival in BMP9/10ib neonates. BMP9/10ib pups develop anemia around P9 and start dying afterward due to widespread vascular and cardiac developmental defects (*19*). A dose of 1 mg/kg DIAG723 (administered s.c. on P3 and P6) significantly increased red blood cell (RBC) counts and fully corrected hemoglobin and hematocrit levels in anemic P9 BMP9/10ib pups (Figs. 6F, 6G). Notably, 1 mg/kg DIAG723 given s.c. on P3, P6, P9, and P12 fully protected pups for up to 20 days against early lethality of the BMP9/10ib pups (Figs. 6H, 6I). Finally, DIAG723 dose-dependently enhanced Smad1/5/8 signaling in HHT2 patient BOECs (Fig. S5), reaffirming our initial finding with DIAG100 that Smad1/5/8 signaling can be activated in the presence of ALK1 LoF mutations in either the extracellular or kinase domain.

**Figure 6:**
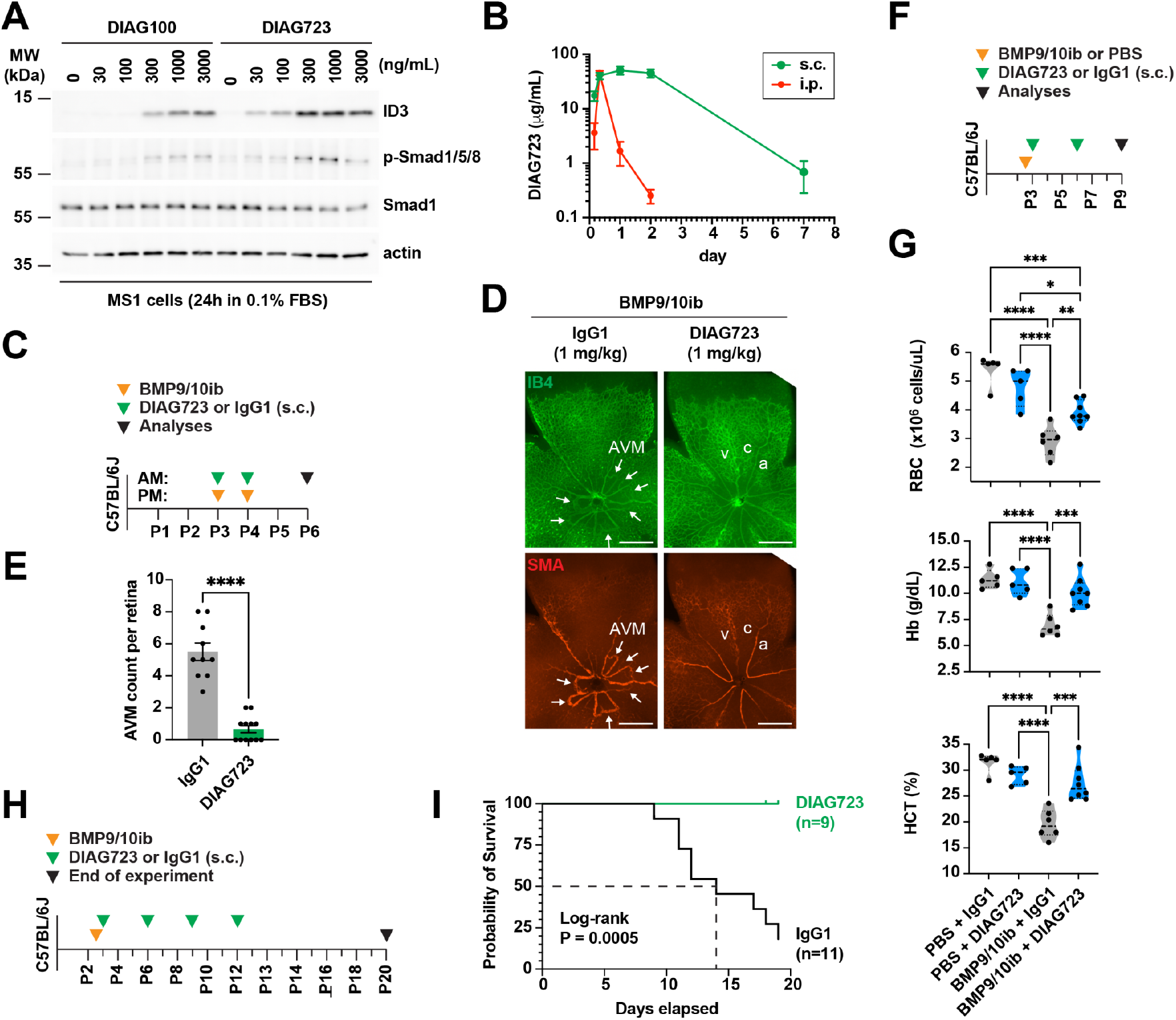
DIAG723 blocks AVM development, anemia, and lethality in BMP9/10ib mice. (**A**) WB analyses of the indicated proteins from MS1 cells treated with different concentrations of DIAG100 and DIAG723 in serum-depleted medium (0.1% FBS) for 24h. (**B**) PK analysis measuring DIAG723 concentration at different time points in the serum of C57BL/6 mice injected (i.p. or s.c.) with 1 mg/kg DIAG723. (**C**) Schematic representation of the injection protocol used in (D, E). (**D**) Representative staining with IB4 (green) and of SMA (red) in whole-mount retinas from BMP9/10ib mice injected (s.c.) with DIAG100 or IgG1 control (1 mg/kg). a, artery; c, capillaries; v, vein. Arrows denote AVMs. Scale bars, 500 µm. (**E**) The bar graph shows the number of retinal AVMs from BMP9/10ib mice treated as in (D). Both eyes of each mouse were analyzed. Data are presented as mean ± s.e.m., unpaired t-test (E). (**F**) Schematic representation of the injection protocol used in (G). (**G**) The violin plots show the distribution of red blood cell (RBC) number, hematocrit (HCT), and hemoglobin (Hb) levels from BMP9/10ib and PBS control mice injected (s.c.) with 1 mg/kg DIAG723 or IgG1 control. Data are presented as mean ± s.e.m., one-way ANOVA with Tukey’s multiple comparisons test. (**H**) Schematic representation of the injection protocol used in (I). (**I**) Kaplan-Meier survival curves and log-rank test of BMP9/10ib mice injected (s.c.) with 1 mg/kg DIAG723 or IgG1 control. For long-term studies in (F-I), BMP9/10 blocking antibodies were injected into the lactating dams. *p< 0.05, **p < 0.01, ***p < 0.001, ****p < 0.0001 (E, G).

## Discussion

Agonist antibodies designed to restore defective signaling pathways show promise as powerful therapeutics for various diseases caused by LoF mutations in receptor complexes (*28*). Here, we provide the first preclinical characterization of BsAbs targeting the extracellular domains of ALK1 and BMPRII (DIAG100 and DIAG723). We show that these BsAbs bound to ALK1 and BMPRII and promoted their proximity association *in vitro* and *in vivo* to activate Smad1/5/8 signaling in ECs (Figs. 1 and 2). *In vivo*, in the BMP9/10ib neonatal retina, we report that ALK1-BMPRII BsAbs interacted with the AVM endothelium (Fig. 2), and dose-dependently prevented AVMs (Figs. 3 and S4). Notably, in this model, DIAG100, after only 48h of treatment, was also able to induce AVM resolution, demonstrating that AVMs can rapidly remodel when ALK1-BMPRII receptor function is restored (Fig. 3). BsAbs could therefore both prevent and treat AVMs. DIAG723, an improved version of DIAG100 with enhanced receptor association and activation properties (Figs. 1 and 6), not only prevented retinal AVMs but also effectively blocked the anemia and early lethality of the BMP9/10ib mice (Fig. 6).

Although retinal vascular manifestations in HHT patients are subtle, new ophthalmic imaging methods now suggest that retinal telangiectasias are common in this population (*29*). In this context, the extensive data produced in the field from the neonatal retinal AVM model are pertinent to the disease process (*30*). Nonetheless, it is important to confirm these findings in other vascular beds and, ideally, in older animals that are not undergoing neonatal angiogenesis. In addition, most HHT cases (∼90%) result from mutations in either *ENG* or *ACVRL1* (*31*). Therefore, aside from the BMP9/10-deficient model (BMP9/10ib mice), we focused our investigation on *Eng*^iECKO^ mice, a model of HHT1. We demonstrated that DIAG100 effectively blocked AVMs in the pubic symphysis and kneecap, as well as improved cardiomegaly in 8-week-old *Eng*^iECKO^ mice (Fig. 3). A popular HHT2 model is the *Acvrl1*^iECKO^ mouse, which develops systemic AVMs during angiogenesis (*32*). Since ALK1 is deleted in this model, testing BsAbs without one of their targets was not feasible. Therefore, we chose an alternative approach by using a KI mouse carrying the HHT2 ALK1-R479* mutation, a model we introduce here for the first time. Since ALK1-R479* KI mice do not spontaneously develop AVMs, at least during the neonatal stage, they were challenged with BMP9/10ib to induce vascular pathology. We found that DIAG100 decreased the number of AVMs in this model, an effect supported by the ALK1 activation properties of the BsAbs in HHT2 patient BOECs, demonstrating that the BsAbs can activate ALK1 signaling in the presence of HHT mutations in ALK1 (Fig. 4). Overall, our results show that forced ALK1-BMPRII agonism restores ALK1 function in different HHT models, thereby preventing AVMs and their associated sequelae such as cardiomegaly, anemia, and early lethality.

AVMs in HHT develop through a multi-hit process that may involve a proangiogenic challenge and complete LoF of an HHT gene (*33*–*35*). In addition to inherited haploinsufficiency in one of the HHT genes, a focal second hit in the form of somatic mutations leading to biallelic LoF in *ACVRL1* or *ENG* has been identified in some ECs of HHT patients’ AVMs (*36, 37*). While it is well known that inherited germline mutations in ECs contribute to disease development, the role of ECs carrying biallelic HHT gene mutations in the development and expansion of the entire AVM remains uncertain (*33*). Studies in mosaic *Acvrl1*^iECKO^ and *Eng*^iECKO^ models have shown that within AVMs, proliferation of wild-type ECs can increase due to neighboring ECs with biallelic *Acvrl1* or *Eng* LoF (*38, 39*). This data suggests that although AVM ECs with somatic HHT gene mutations represent a small portion of all the ECs forming AVMs, a spreading mechanism originating from these cells may contribute to the disease process. Our results show that clustering ALK1 and BMPRII with a BsAb can effectively prevent AVM pathology despite ENG deficiency in *Eng*^iECKO^ mice (Fig. 3), demonstrating the potential of this method to treat ECs with biallelic *ENG* LoF. When considering biallelic *ACVRL1* LoF mutations (more specifically, those affecting the kinase domain), an ALK1-BMPRII agonism strategy might be ineffective in the ECs that carry these mutations. However, this strategy could still potentially rescue the adjacent AVM ECs that only carry a germline ALK1 mutation (per our observations, Fig. 4). To address this, more advanced models with mosaic deletion of *Acvrl1—*preferably in an HHT2 germline mutant background—will need to be developed.

Our transcriptomic analysis reveals that ALK1-BMPRII activation by DIAG100 controls a specific subset of BMP9-regulated genes without triggering an off-target transcriptional response (Fig. 5). Within this gene subset, we identified genes modulated with similar amplitude by DIAG100 and BMP9 (including ID1 and ID3) and genes responding more weakly to DIAG100 than to BMP9 (including SGK1). These results are significant because they show that the BsAb did not trigger obvious off-target transcriptional responses, as it did not engage pathways or targets outside BMP9 control. Additionally, this data identifies a subset of genes that are similarly regulated by DIAG100 and BMP9 and may serve as more direct factors in a signaling response that maintains vascular quiescence and prevents AVMs. Further research will be necessary to determine whether this gene subset is part of a specific hub involved in a particular cellular defect or mechanism of AVM development.

Currently, no therapies have been approved for treating HHT. The blood vessel walls of AVMs in HHT are fragile and prone to rupture, leading to chronic bleeding, acute hemorrhage, and complications from shunting through the vascular malformations. As a result, approximately 60% of HHT patients develop anemia and require ongoing iron infusions and/or RBC transfusions, making HHT a resource-intensive and very costly rare disease (*5, 6, 40*). Current treatments mainly focus on managing symptoms, such as coagulation therapy, embolization, cautery, and various surgical procedures. Some therapies are under development and primarily target the VEGFR2-PI3K-Akt-mTOR pathway or aim to reduce bleeding (*14, 41*). However, none address the underlying cause of the disease, which involves a destabilized vessel wall due to ENG-ALK1 receptor LoF. As described, an ALK1-BMPRII agonism achieved via clustering of the two receptors with a bispecific ALK1:BMPRII antibody restores deficient signaling caused by ENG-ALK1 receptor LoF. Hence, this BsAb approach represents a novel, potentially disease-modifying strategy to treat the underlying cause of HHT. Beyond its potential to correct HHT, the bispecific clustering of ALK1 and BMPRII might also have therapeutic potential for pulmonary arterial hypertension, a condition associated with defective ALK1-BMPRII signaling and LoF mutations in BMPRII, which have also been reported in HHT patients. In conclusion, this study highlights the therapeutic potential of a BsAb-driven ALK1-BMPRII agonism approach in HHT.

## Materials and Methods

Materials and Methods are available as supplementary materials.

## Acknowledgements

We thank Dr. Christine N. Metz (Feinstein Institutes for Medical Research) for generously providing us with HUVECs, and Dr. Peter J. Romanienko from the Genome Editing Shared Resource at Rutgers Cancer Institute (New Brunswick, NJ, USA) for generating the ALK1-R479* KI mice. We also thank Dr. Jean-Christophe Hus (Diagonal Therapeutics, Watertown, MA, USA) for his scientific insights on biologics discovery.

## Accession Numbers

RNA-seq reads were deposited in the European Nucleotide Archive (ENA): Accession # PRJEB95834 / ERP178587.

## Author contributions

P.M., P.A., and S.Q. conceived the study. S.Q., H.Z., Z.W., A.S., M.G., R.K., H.T., H.Y., X.W., and S.G. developed the methods and conducted the experiments. H.N.S., M.B., and E.C. recruited donors and collected blood samples. F.C. carried out the bioinformatic analyses. H.M.A. provided material. P.M., S.Q., P.A., A.S., M.G., and H.M.A. wrote, reviewed, or edited the manuscript.

