## Supplementary Material for "ALK1-BMPRII agonism by clustering bispecific antibodies treats hereditary hemorrhagic telangiectasia"

### Materials and Methods

#### *Mice and treatments*

All animal procedures were performed in accordance with protocols approved by The Feinstein Institutes for Medical Research Institutional Animal Care and Use Committee and conformed to the National Institutes of Health (NIH) Guide for the Care and Use of Laboratory Animals and ARRIVE guidelines. Animals were housed under a 12h light/dark cycle at temperatures of 18–23$^{o}$C with 40-60% humidity and ad libitum access to food and water. C57BL/6 mice from The Jackson Laboratory were used for the BMP9/10ib studies. Anti-BMP9 (12.5 mg/kg, R&D Systems, MAB3209) and anti-BMP10 (25 mg/kg, R&D Systems, MAB2926) antibodies were administered by i.p. injection into pups on P3 and P4 [as described in Ref. (*20*)] or into lactating dams on P2.5 [as before (*18*)]. Mice were sacrificed on P6, P7, or P9, depending on the analysis. *Eng*$^{f/f}$ (*21*, *22*) were crossed with *Cdh5*-CreER$^{T2}$ mice [Taconic, 13073; (*42*)] to generate EC-specific, tamoxifen-inducible *Eng* knockout (*Eng*$^{iECKO}$) mice. *Eng*$^{iECKO}$ mice were also crossed with *Rosa26*-tdTomato reporter mice (Ai14 strain; The Jackson Laboratory, 007914) to assess recombination efficiency. Gene deletion was induced in 6-week-old *Eng*$^{iECKO}$ female mice by intragastric administration of 2 mg tamoxifen (Sigma, T5648) on day 0 (D0), D1, and D2. The generation of the ALK1-R479* KI mice was outsourced to the Genome Editing Shared Resource at Rutgers Cancer Institute, New Brunswick, NJ, USA. ALK1-R479* (human equivalent of mouse ALK1-R478* mutant) KI mice were generated using CRISPR-Cas9. C57BL/6 embryos were microinjected with a mixture containing Cas9 protein (IDT), an sgRNA (Millipore Sigma), and an ssODN (IDT), which contained homology arms and the R478* mutation. The sgRNA used was TCTTTATGCGCAGTGCGGTGAGG (PAM is underlined), and the donor oligo sequence was GTCCTCTCCGGGCTGGCCCAGATGATGAGAGAGTGCTGGTACCCCAACCCCTCTGCTtGaCTCACCGCACTGCGCATAAAGAAGACATTGCAGAAGCTCAGTCACAATCCAGAGAAGCCCAAAGTG (R478* sequence change is shown in lower case; the change modified the PAM sequence to avoid recutting). Founders were screened by PCR and digested with the restriction enzyme HinfI (introduced by R478* and PAM change). NGS sequencing (Azenta Life

Sciences) was used to confirm the R478* change. Confirmed founders were bred, and progeny were screened using PCR primers ACVRL1A (CTGCTATGTCTCCCGATCCTGAG) and ACVRL1B (CTCAGCTGTATTTTTGGCTGGATG) for the gain of HinfI restriction site and by Sanger sequencing of PCR products. ALK1-R479* KI mice were injected with BMP9/10 blocking antibodies (as described above) to generate BMP9/10ib;*Acvrl1*$^{R479*/+}$ mice. The sex of the pups was not determined. At the indicated times, mice were administered i.p. or s.c. injections of DIAG100, DIAG723, or human IgG1 control in PBS.

***Antibody production***

Antibodies targeting ALK1 and BMPRII were identified from a variety of sources, including both immune and synthetic libraries. Antibodies were transiently transfected using the Expi293 system according to the manufacturer's instructions (ThermoFisher Scientific). Cells were harvested six days post-transfection and harvested using batch purification with MabSelect resin (Cytiva). Purity of the final product was assessed using SDS-PAGE and analytical gel filtration. Individual antibodies were screened for affinity using ELISA and SPR and then tested for agonistic activity in high-throughput formats using the DiscoverX assay. Promising binder pairs were further optimized in regard to format, linkers, and valency using the DiscoverX assay.

***Affinity assays***

Binding affinities were measured using the Carterra LSA equipped with an HC30M chip. Binding assays were carried out at 25°C with HBSTE buffer (10 mM HEPES, pH 7.4, 150 mM NaCl, 3 mM EDTA, 0.05% Tween-20) supplemented with 0.5 g/L BSA. To prepare the lawn, the chip was activated with 40 mM EDC and 10 mM S-NHS in 100 mM MES (pH 5.5). Coupling of a goat anti-human IgG antibody (Southern Biotech, 2087-01) with standard immobilization was done for 15 minutes in 10 mM sodium acetate (pH 4.5), quenched using 1M ethanolamine, and washed with 10 mM glycine (pH 2.0). The anti-human IgG lawn surface was then used to capture a panel of selected antibodies at 10 μg/mL, 2 μg/mL, or 400 ng/mL. Monomeric human ALK1, murine ALK1, or human BMPRII was injected over the captured antibody array at 6 concentrations in a 3-fold dilution series starting at 500 nM. Binding data was referenced and blanked from the buffer, then globally fit to a 1:1 Langmuir binding model for estimation of ka, kd, and KD using Carterra Kinetics Software.

***DiscoverX assay***

The PathHunter™ U2OS ACVRL1/BMPR2 Dimerization cell line (Eurofins, 93-0962C3) was used as described. Briefly, confluent cells were detached (Eurofins, 92-0009) and spun down at 300xg for 5 min. Cells were resuspended in plating reagent (Eurofins, 93-0563R22A) at a density of $2.5x10^5$ cells/mL, and 20 µL was added to each well of a 384-well assay plate. The plate was incubated overnight at 37°C, 5% $CO_2$. Antibodies and the BMP9 control (R&D Systems, 3209-BP/CF) were diluted 10-fold from 100 nM with assay dilution buffer (Eurofins, 92-0023). 5 µL of the diluted sample was added to each well of the plate. The plate was then incubated at 37°C for 3h. 25 µL of freshly prepared detection reagent (Eurofins, 93-0247) was added to each well and incubated at room temperature for 1h in the dark. A VarioSkan LUX plate reader was used to detect the chemiluminescence signal.

***Cell cultures***

MS1 cells (ATCC, CRL-2279) were grown in Dulbecco's modified Eagle medium (DMEM; ThermoFisher Scientific, 11965118), 10% fetal bovine serum (FBS; Cytiva, SH30070.03), and 1% penicillin-streptomycin (Gibco, 15140122) until they reached approximately 90% confluence and then were incubated in fresh complete medium or serum-depleted medium (0.1% FBS) before treatments. HUVECs were isolated from umbilical veins obtained from anonymous donors (*43*) and were cultured in EC medium (ECM; ScienCell, 1001) supplemented with 5% FBS and EC growth supplement (ScienCell, 1052). For bulk RNA-seq, HUVECs were grown in 6-well plates using ECM until they reached approximately 90% confluence and then were incubated in serum-depleted medium (0.5% FBS without supplement) for 2h and treated for 4h with 1 µg/mL IgG1, 1 µg/mL DIAG100, or 1 ng/mL human rBMP9 (R&D Systems, 3209-BP). BOECs were isolated from blood samples taken from HHT patients during routine visits at the Northwell Health Otolaryngology & Facial Plastics division, following the previously described protocol (*15*). BOECs were cultured in EBM-2 basal medium (Lonza, CC-3156) supplemented with 10% FBS and EGM-2 MV SingleQuots supplement pack (Lonza, CC-4147). HMEC-1 (CRL-3243), TIME cells (CRL-4025), and HPAEC (PCS-100-022) were obtained from ATCC and cultured following ATCC's recommended protocols.

***p-SMAD1 ELISA***

HMEC-1, TIME cells, and HPAECs were plated in growth media at 20,000 cells/well in a 96-well plate overnight, then starved in basal media for 4h before treatment for 1h with DIAG723, DIAG100, or rBMP9 (R&D Systems, 3209-BP/CF). After treatment, cells were washed with PBS, and p-SMAD1 levels were measured using the SMAD1 (pS463/465) ELISA Kit (Abcam, AB186036) and following the manufacturer's protocol. A relative standard curve for the assay was built by making 1:2 dilutions of the control sample included in the kit, where the undiluted control is assigned a value of 100. Sample values were calculated relative to the standard curve.

***Western blot analyses***

Cells were solubilized in RIPA buffer (EMD Millipore, 20-188) supplemented with 1× Complete protease inhibitor cocktail (Roche, 11697498001). 5-20 μg of protein (depending on the primary antibody) was separated by SDS-PAGE and transferred onto nitrocellulose membranes, which were probed with primary and secondary antibodies. A standard enhanced chemiluminescence detection procedure was used. Primary antibodies against actin (1:5,000; BD Biosciences, 612656), ID1 (1:1,000; BioCheck, BCH1/195-14), ID3 (1:1,000; BioCheck, BCH-4/#17-3), p-Smad1/5/8 (1:1,000, Cell Signaling Technology, 13820), Smad1 (1:1,000, Cell Signaling Technology, 6944), and SGK1 (1:1,000, Cell Signaling Technology, 12103) were used.

***Antibody intracardiac injection***

DIAG100 conjugation to Alexa Fluor 647 (DIAG100-AF647) was achieved via amine-reactive N-hydroxysuccinimide (NHS) ester chemistry, which targets primary amines predominantly found on lysine residues. Briefly, DIAG100 was incubated with up to 10 molar excess of Alexa Fluor 647 NHS ester (ThermoFisher Scientific, A20106) reconstituted in anhydrous DMSO (dimethyl sulfoxide) for 1h at room temperature. Conjugated DIAG100 was separated from free dye using a Zeba Spin desalting column (ThermoFisher Scientific) and confirmed for purity (> 90%) using HPLC-SEC before aliquoting and deep-freezing for subsequent use. The ratio of Alexa Fluor 647 dye molecule to antibody was estimated to be 2.31 using the Beer-Lambert law with the measured absorbance of the antibody (A280) and dye (A650). DIAG100-AF647, human IgG1-AF647 (DDXCH01A647-100, Novus Biologicals), and CD31 Vio Bright B515

(CD31-B515; Miltenyi Biotec, 130-111-358) antibodies were injected intracardially in P7 neonates under anaesthesia (deep hypothermia). The thoracic cavity was opened, and 50 µL of antibody mixture was injected with a syringe (31G needle) directly into the left ventricle of the heart, over approximately 30 sec. The heart continued to beat for approximately 5 min following the injection. The pups were then perfused through the left ventricle with 10 mL HBSS (Gibco, 2276886) supplemented with 100 IU/ml heparin (EMD Millipore, 375095-500KU), followed by 5 mL of 4% paraformaldehyde (Electron Microscopy Sciences, 19208) in PBS (Cytiva, SH30378.02), using a BS-300 syringe pump (Braintree Scientific, inc.) connected to a 27G needle Scalo Vein set, set at a pumping rate of 0.6 mL/min. Retinas were harvested, processed, and directly imaged as described below.

***Retinal immunofluorescence***

Retinas were processed as previously described (*18*, *20*). In brief, after the mice were killed, their eyes were dissected and then completely immersed in ice-cold 4% paraformaldehyde in PBS for 35 min. The eyes were transferred to ice-cold PBS for dissection. The cornea, lens, and pigmented layers were removed from the retinas, and four incisions were made with Vannas spring scissors (Fine Science Tools, 91500-09) for flat mounting. PBS was removed, and retinas were immersed in 100% ice-cold methanol for 15 min or stored at -20ºC. Retinas were permeabilized with PBS containing 0.3% Triton X-100 for 15 min, and immersed in a PBS blocking solution containing 10% heat-inactivated goat serum (ThermoFisher Scientific, 16210072), 0.3% Triton X-100, and 0.2% BSA (HIS-PBST) for 1h at room temperature. Retinas were then stained at 4°C overnight with isolectin GS-IB4 conjugated to Alexa Fluor 488 (1:300 in HIS-PBST; ThermoFisher Scientific, I21411) and anti-α smooth muscle actin-Cy3 antibody (1:200 dilution; Sigma-Aldrich, C6198). Retinas were then washed three times with HIS-PBST. Retinas were mounted on slides using an antifade mountant (ThermoFisher Scientific, P36965) and sealed with nail polish. Retinas were imaged using a Zeiss LSM 900 confocal microscope and a ThermoFisher EVOS M7000 microscope.

***ALK1-BMPRII Proximal ligation assay (PLA)***

PLA was performed on paraformaldehyde-fixed and permeabilized mouse retinas as per the manufacturer's instructions (Sigma, DUO92102; DUO92004; DUO92002) using rabbit anti-ALK1 (1:100; GeneTex, GTX100035) and mouse anti-BMPRII (1:200; GeneTex, GTX60415) antibodies, and costained with Alexa Fluor 488-conjugated IB4. Quantification was performed on single focal planes by determining the total number of PLA signals per IB4-positive area in the arteries proximal to the optic nerve center.

#### *Blue latex dye perfusion*

Six to eight-week-old female mice were perfused with blue latex dye (Connecticut Valley Biological Supply, BR80B) under terminal anesthesia. Prior to the latex blue injection, mice received an i.p. injection of 0.3 mL of 500 IU/mL heparin in HBSS. Then, they were anesthetized with 100 μL of ketamine/xylazine solution [100 mg/kg ketamine (Covetrus, DC:11695-0703-1), 10 mg/kg xylazine (Covetrus, NDC:11695-4024-1)]. The chest was opened to expose the heart and lungs, and a PE50 tube was inserted downward into the descending aorta and advanced toward the diaphragm. The right atrium was incised to facilitate drainage of blood and solution, and the blood was washed out by perfusing 10 mL of pre-warmed HBSS supplemented with 100 IU/mL heparin using a BS-300 syringe pump. Then, 0.3 mL blue latex dye was injected. Tissues were left *in situ* for ~15 min at room temperature. The pubic symphysis was then dissected and imaged using an Olympus SZX7 stereomicroscope attached to an Olympus DP27 camera. Measurements and quantifications were performed using Fiji.

#### *RNA Sequencing and Analysis*

Frozen HUVEC pellets were sent to Azenta/GENEWIZ for total RNA extraction, library preparation, and sequencing. Bulk RNA sequencing was performed using the Illumina platform, following standard procedures (2x150 bp, ~20 million paired-end reads per sample). Data analysis, including quality control, alignment, and differential gene expression analysis, was conducted by Azenta/GENEWIZ bioinformatics specialists using their standard pipeline. Venny2.1 was utilized to create the Venn diagrams. Gene expression data were filtered by gene list and expressed as a Z-score by row (Z-score calculated across the expression values of each gene), then rendered as a heatmap with the Python Seaborn *clustermap* function (clustering by row/gene was enabled).

**Supplementary Figures**

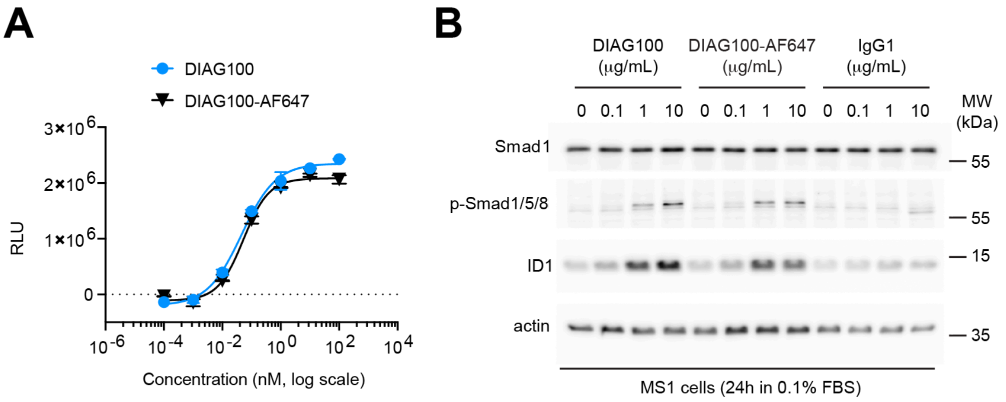


**Figure S1: DIAG100-AF647 retains its clustering capabilities and Smad1/5/8 signaling activation properties.** (**A**) DiscoverX assay measuring ALK1 and BMPRII proximity association by DIAG100 and DIAG100-AF647. (**B**) WB analyses of the indicated proteins from MS1 cells treated with different concentrations of DIAG100, DIAG100-AF647, or IgG1 control in serum-depleted medium (0.1% FBS) for 24h.

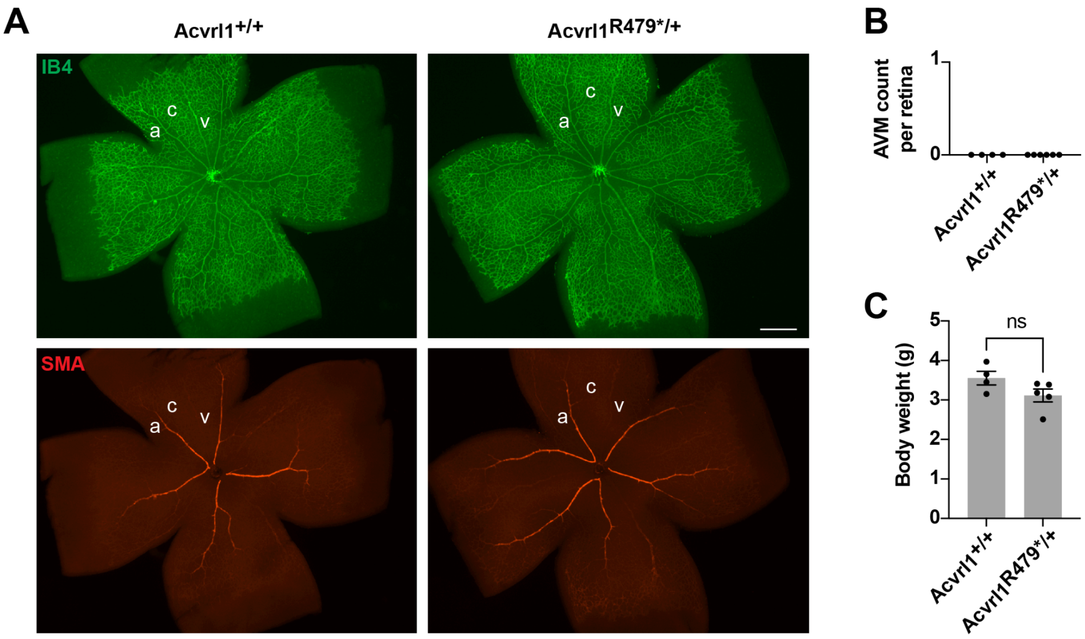


**Figure S2: *Acvrl1*$^{R479*/+}$ mice develop no obvious vascular defects in the retina at the neonatal stage.** (**A**) Representative staining with IB4 (green) and of SMA (red) in whole-mount retinas from P6 *Acvrl1*$^{R479*/+}$ mice and *Acvrl1*$^{+/+}$ control littermates. (**B, C**) The bar graphs show the number of retinal AVMs (B) and body weight (C) of mice as in (A). Data are presented as mean ± s.e.m.; unpaired t-test in (C). Both eyes of each mouse were analyzed.

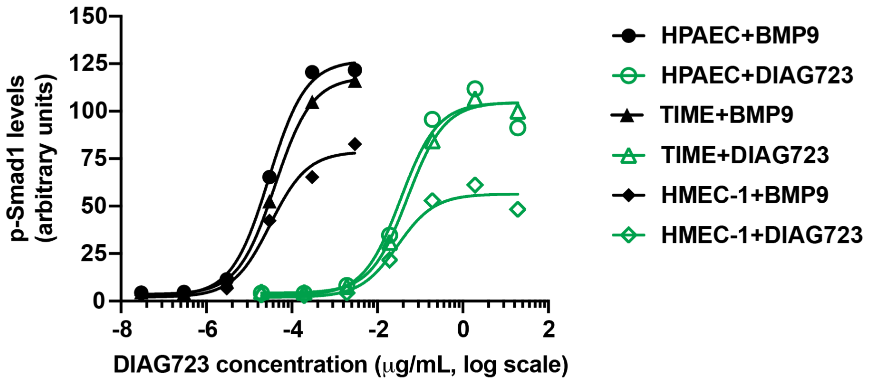


**Figure S3: Smad1 signaling activity of DIAG723 in HPAECs, TIME cells, and HMEC-1.** p-Smad1 level measurements from the indicated human endothelial cells starved in basal media for 4h and then treated with different concentrations of DIAG723 for 1h.

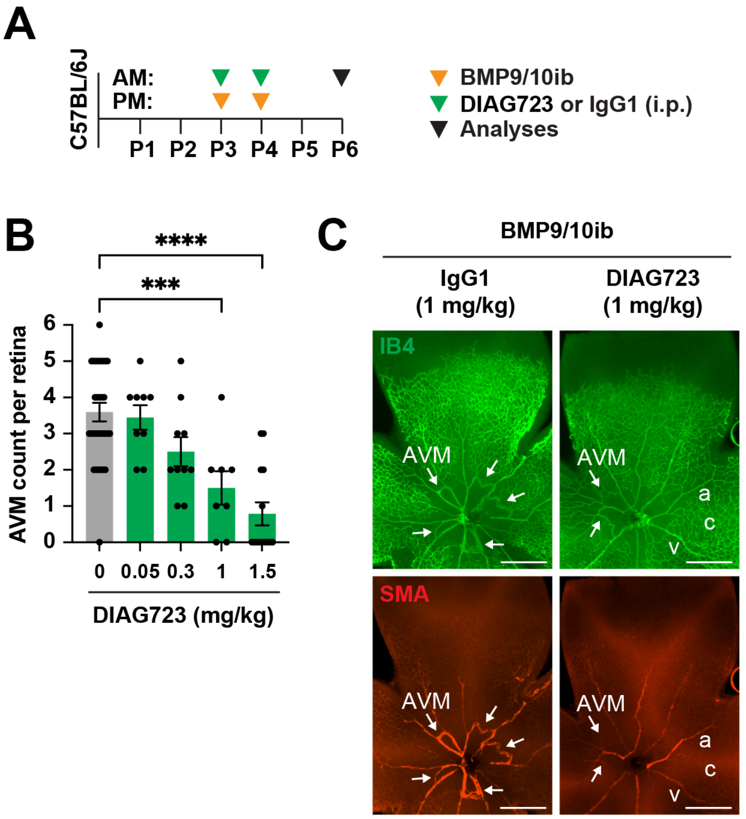


**Figure S4: DIAG723 prevents AVM development in BMP9/10ib mice.** (**A**) Schematic representation of the injection protocol used in (B) and (C). (**B**) The bar graph shows the number of retinal AVMs from BMP9/10ib mice treated with the indicated doses of DIAG723. Data are presented as mean ± s.e.m., one-way ANOVA with Tukey's multiple comparisons test. Both eyes of each mouse were analyzed. (**C**) Representative staining with IB4 (green) and of SMA (red) in whole-mount retinas from BMP9/10ib mice treated (i.p.) with DIAG723 or IgG1 control (1 mg/kg).

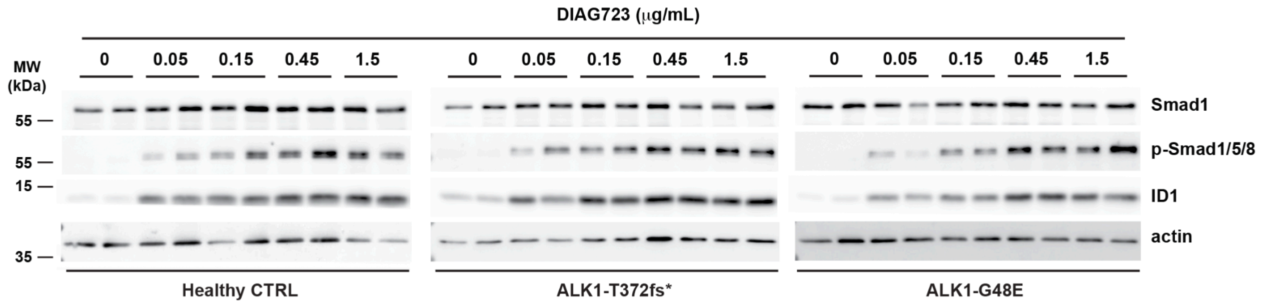


**Figure S5: DIAG723 activates ALK1 signaling in HHT2 patient BOECs.** (**A**) WB analyses of the indicated proteins from healthy donor and HHT2 patient BOECs serum-depleted for 3h (0.1% FBS) and then treated with the indicated concentrations of DIAG723 for 2h.
